# Systemic induced plant resistance genes expression in winter wheat under treatment with antagonistic bacteria of *Fusarium sp*

**DOI:** 10.1101/2025.10.16.682632

**Authors:** Kamilla Mekhantseva, Nikita Vasilchenko, Evgeniya Prazdnova, Alexander Usatov, Maria Mazanko, Vladimir Chistyakov

## Abstract

**Introduction:** Fusarium species are devastating wheat pathogens. Plant growth-promoting rhizobacteria (PGPR) offer an effective biocontrol alternative. This study investigated the capacity of a Bacillus and Paenibacillus consortium to induce systemic resistance in winter wheat.

**Methods:** Winter wheat plants were treated with a ten-strain PGPR consortium, with or without Fusarium oxysporum and F. graminearum. Expression of five PR-gene families was quantified in roots and shoots at 7 and 14 days post-treatment via RT-qPCR. Morphometric parameters were tracked for 28 days.

**Results:** The consortium elicited a potent defense response. In roots at 7 days, PR-1 and PR-6 expression surged 16-fold and 38-fold, respectively. Defense activation in shoots was delayed, peaking at 14 days. After 28 days, PGPR-treated plants exhibited an 8% increase in shoot length.

**Discussion:** The results confirm that the bacterial strains effectively prime the wheat immune system, modulating early defense pathways and enhancing plant growth.

## 1. Introduction

Cereal crops form the basis of the diet of the population in many countries [1], with wheat being one of the top five most produced cereal grains. In Russia, wheat occupies about 70% of the sown area. Wheat yield is affected by abiotic and biotic stresses. Among biotic stresses, fungal exposure, including fungi of the genus *Fusarium*, are major concerns. In particular, *Fusarium* exposure might result in seedling death and deterioration of crop health, leading to reduced yields [2], loss of quality and marketability [3], as well as accumulation of mycotoxins rendering the grain hazardous to mammals.

Of the 30 different diseases that affect crops, Fusarium head blight and Fusarium wilt are considered the most serious. Outbreaks of these diseases occur every fourth to fifth year in the United States, China, European Union, the United Kingdom, Africa, Brazil and other countries. Fusarium wilt is caused by *Fusarium* fungi, with *Fusarium graminearum* and *Fusarium oxysporum* being some of the most prevalent species. They affect the plant’s vascular system [4] and spread through the soil via highly resilient spores, which can persist in the soil for decades [5]. While Fusarium spp. are often pathogenic, their presence alone does not guarantee disease; plant responses vary by strain, environment, and host immunity.

For a long time, chemical fungicides were primary means of *Fusarium* controlling. However, they have several drawbacks such as high cost; negative environmental impacts as fungicides accumulate in the soil; limited period of effectiveness due to a rapid adaptation of fungi to new chemical agents. Currently, one of the most effective solutions of this problem involves using microorganisms, particularly, plant growth-promoting rhizobacteria (PGPR) as biocontrol agents [1, 6]. They may mitigate *Fusarium*-associated stress by priming plant immunity. These bacteria form beneficial relationships with plants and may enhance growth and yields [7]. Winter wheat has complex defense mechanisms against *Fusarium*, some of which involve the synthesis of PR (pathogenesis-related) proteins. The expression of *PR*-genes in wheat can increase up to 70-fold during *Fusarium* exposure [8]. However, the effect of PGPR on *PR*-genes expression in wheat under *Fusarium* exposure is not yet well understood. We hypothesize that PGPR may influence *PR*-gene expression, representing an additional mechanism of action of rhizosphere strains antagonistic to *Fusarium*. This effect could potentially be mediated by the same metabolites that are responsible for antagonism against fungi.

Bacterial preparations can suppress Fusarium wilt in cultivated plants, such as, for example, “Orgamika F” for cabbage [9] or a “Code of Balance” biopreparation tested on wheat [10]. For optimal use of biocontrol agents, it is crucial to understand their effects on plants. In particular, their ability to induce systemic resistance is key to both application and fundamental research. Biocontrol agents that have such ability show a broader range of applications, such as selecting specific antagonists for specific strains of the fungi.

This study examines early-stage interactions — focusing on PR-gene expression and growth trends— in wheat exposed to *Fusarium* and PGPR. The aim of this study was to investigate the effect of a biocontrol preparation, consisting of 10 potential PGPR strains antagonistic to *Fusarium*, on the expression of winter wheat immunity genes and plant growth parameters in a model experiment.

## 2. Materials and methods

### 2.1. Plant, bacterial and fungal species

Seeds of winter wheat (*Triticum aestivum L*.) of “Tatiana” variety (developed in Scientific and Production Center of Grain Farming named after A.I. Baraev, Kazakhstan Republic) were used in this work. Conidia of *Fusarium oxysporum* and *Fusarium graminearum* species (standard strain from the Southern Federal University’s microbiological collection, obtained from agricultural field in Krasnodar region, Russia) were used for exposure, because these strains have been shown to cause plant diseases in the fields from which they were isolated [11].

Eight strains of bacteria of the genus *Paenibacillus* and 2 strains of the genus *Bacillus* were used for bacterial treatment. All these strains were isolated from soils of fields in Krasnodar Region, Russia. In previous studies, they demonstrated antifungal activity against fungi of the genus *Fusarium* both in laboratory conditions and in field experiments [12, 13].This preparation is patented in Russia under the name «Code of balance F1», and the genomes of these bacteria are deposited in NCBI database (*ID 806727 - BioProject - NCBI*) (Table 1).

**Table 1.** Bacterial strains of the Code of balance F1 biocontrol agent.

| Strains | Genbank accession number |
| --- | --- |
| *Paenibacillus jamilae* K 1.14 | MN219707 (16 S) |
| *Paenibacillus peoriae* O 1.27 | MN219728 (16 S) |
| *Paenibacillus peoriae* O 2.11 | MN220137 (16 S) |
| *Paenibacillus peoriae* R 3.13 | MN220147 (16 S) |
| *Paenibacillus peoriae* R 4.5 | MN220138 (16 S) |
| *Bacillus amyloliquefaciens* R 4.6 | MN220139 (16 S) |
| *Paenibacillus jamilae* R 4.24 | MN220140 (16 S) |
| *Paenibacillus polymyxa* R 5.31 | MN220144 (16 S) |
| *Paenibacillus peoriae* R 6.14 | MN220145 (16 S) |
| *Bacillus amyloliquefaciens* V 3.14 | MN220146 (16 S) |

Table 1 - Bacterial strains of the Code of balance F1 biocontrol agent
| **Strains** | **Genbank accession number** |
| --- | --- |
| *Paenibacillus jamilae* K 1.14 | MN219707 (16 S) |
| *Paenibacillus peoriae* O 1.27 | MN219728 (16 S) |
| *Paenibacillus peoriae* O 2.11 | MN220137 (16 S) |
| *Paenibacillus peoriae* R 3.13 | MN220147 (16 S) |
| *Paenibacillus peoriae* R 4.5 | MN220138 (16 S) |
| *Bacillus amyloliquefaciens* R 4.6 | MN220139 (16 S) |
| *Paenibacillus jamilae* R 4.24 | MN220140 (16 S) |
| *Paenibacillus polymyxa* R 5.31 | MN220144 (16 S) |
| *Paenibacillus peoriae* R 6.14 | MN220145 (16 S) |
| *Bacillus amyloliquefaciens* V 3.14 | MN220146 (16 S) |

Strains were isolated from rhizosphere soil (at a depth of 0-10 cm in close proximity to plant roots) sampled on a wheat field in Krasnodar area, Russia. Identification was carried out by 16s RNA.

### 2.2. Inoculation and cultivation of experimental and control wheat groups

Plants were treated with prepared suspension of *Fusarium oxysporum* and *Fusarium* graminearum conidia. Fungi were pre-cultured for 14 days with the use of malt extract agar in a petri dish, then mycelium was washed with physiological solution and adjusted to a concentration of 10^6^ conidia/1 ml with serial dilutions and counting conidia in a Burker’s chamber. To treat winter wheat seeds, they were immersed in the fungal conidia suspension, at a ratio of 1 ml of suspension per 100 g of seeds. The seeds were then removed from the suspension and air-dried. 1 ml was then dissolved in a larger volume of water. Thus, the final content was 5*10^5 CFU/ml, and the volume of water in which the seeds were soaked was 50 ml per 100 g.

For seed treatment with antagonist strains, the bacteria were incubated on solid nutrient medium (potato dextrose agar) for 24 hours, then harvested and resuspended in sterile saline. All strains were added to the final preparation in the same proportion. Seeds were treated at with 50 ml of suspension of 5·106CFU/1 ml for 100 g of seeds.

Seeds soaked in sterile physiological solution were used as a control group.

Fifty seeds for each experimental group were planted per 15-liter pots at a depth of 1-2 cm. The soil nutrient levels (mg/l) were: N (100-180), P2O5 (135-255), K2O (115-215), with a pH of 5-6. Plants were grown indoors at 18-22 °C, with a 12-hour photoperiodartificial light, and periodic watering. Full spectrum Goodland phytolamps were used, and a 2-meter-high polyethylene barrier separated experimental groups to prevent cross-contamination. Humidity was maintained above 70% in all pots. Plant material for further analysis was collected 7 and 14 days after planting in 3 biological replicates.

In all the Figures, four types of treatment are shown: “bacteria” – potential PGPR chosen, “fungi” – *Fusarium* solution, “combination” - potential PGPR and *Fusarium* solution, “control” – no specific treatment.

### 2.3. RNA isolation from wheat plants and gene expression analysis

Gene expression was assessed at 7 and 14 days to see early and intermediate molecular responses to inoculation, consistent with prior studies showing that plant defense genes, including PR genes, are differentially regulated within the first two weeks of fungal exposure [14, 16].

RNA isolation was performed by preliminary shredding of tissues in liquid nitrogen using a porcelain pestle and mortar, followed by extraction with ExtractRNA kit (Eurogen, Russia). The isolated RNA was purified using CleanRNA Standard kit (Eurogen, Russia). cDNA synthesis was performed using MMLV RT Kit (Eurogen, Russia).

### 2.4. RT-PCR and melting curve analysis

Expression of PR-genes was analyzed by quantitative real-time PCR using qPCRmix - HS SYBR kit (Eurogen, Russia) with relative expression level estimation by 2^-ddCt^ method. Primer sequences are presented in Table 2 (Supplementary materials). Each PCR sample contained 10.4 μl of water for PCR, 4 μl of qPCR–mix SYBR, a primer mixture (Forward and Reverse) in a volume of 0.6 μl, 5 μl of cDNA (concentration of 10 ng/μl).

**Table 2.**
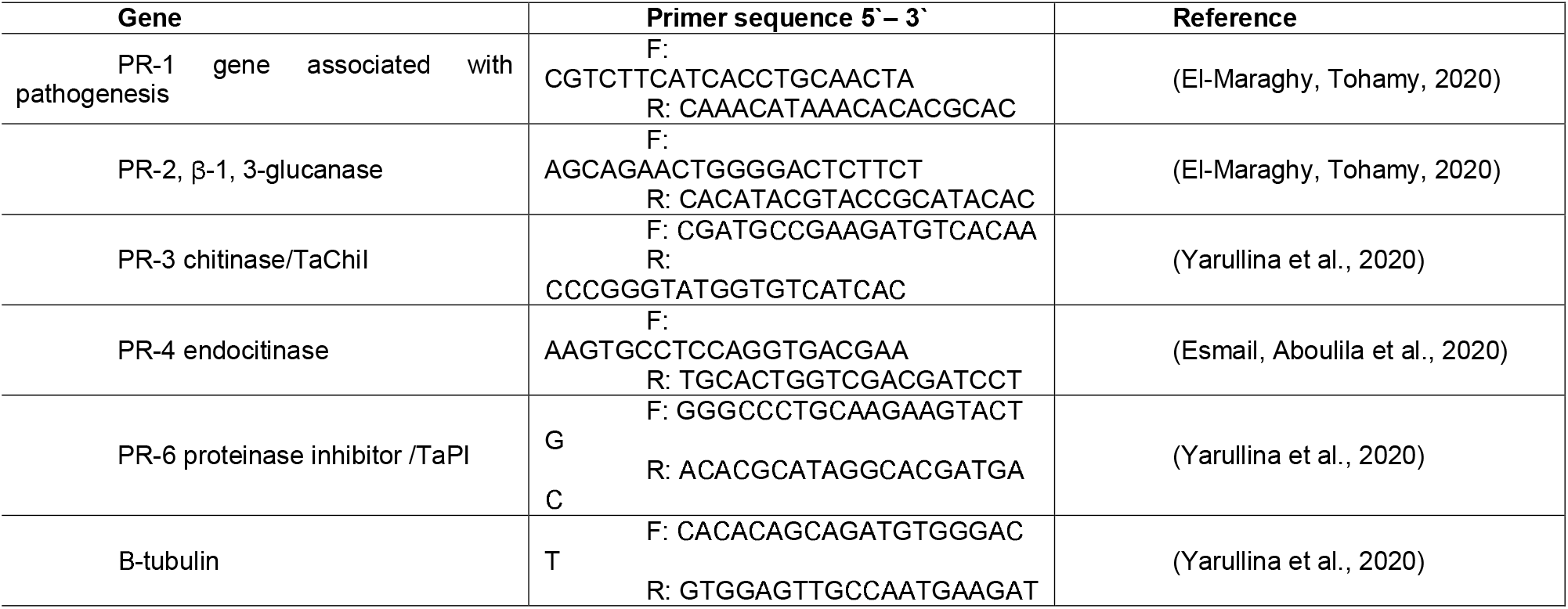
Primers to *PR*-genes and housekeeping genes.

All runs were performed with a QuantStudio 5 (Thermofisher Scientific) using the following program: pre-denaturation at 95 °C for 5 minutes, denaturation at 95 °C for 30 seconds, annealing of primers at 57 °C for 30 seconds, elongation at 72 °C for 30 seconds.

The reaction was performed in 45 cycles. After PCR, melting curves were analyzed using the following parameters: elongation at 72 °C for 15 seconds, maintaining temperature at 55 °C for 1 minute, gradual increase in temperature with a step of 0.2 °C per second, denaturation at 95 °C for 15 seconds.

### 2.5. Housekeeping genes validation

Reference gene stability was evaluated by calculating the coefficient of variation (CV) of Ct values across triplicates. *β-tubulin* and *cdc* were chosen as potential housekeeping genes. *β-tubulin* showed superior stability demonstrating lower CV coefficients than cdc in most groups (Table 6, Supplementary material). Based on comprehensive CV analysis, β-tubulin was selected as the primary reference gene for normalization.

### 2.6. Morphometric measurements

For morphometric evaluation, plant samples were collected at 7, 14, and 28 days after cultivation. At each time point, three plants were randomly selected per pot, carefully extracted to preserve the root system, and rinsed with distilled water to remove substrate residues. The following parameters were measured using a digital caliper (accuracy ±0.05 mm): root length - from the base of the primary root to the tip of the longest lateral root, shoot length - from the stem base to the apex of the longest leaf.

Plants were placed on a flat surface under consistent lighting to ensure measurement accuracy. Each sample was measured once by a single operator. Subsequent measurements were performed on newly selected plants to avoid pseudo-replication.

Fresh weight measurements were conducted immediately after rinsing using a precision laboratory balance, accuracy ±0.01 g). Excess moisture was gently blotted with filter paper before weighing to ensure consistency.

Morphological traits (shoot and root length) were additionally measured for 28 days to study long-term and cumulative effects of exposure and treatment. Also, the 28-day assessment aligns with the complete inoculation cycle of Fusarium species [17].

### 2.7. Statistical Analysis

Prior to conducting comparative analyses, the assumption of normality was evaluated for all groups using the Shapiro-Wilk test (α = 0.05). For datasets satisfying normality assumptions, parametric analysis was performed using one-way analysis of variance (ANOVA) to detect overall differences among treatment groups. When ANOVA indicated significant effects (p < 0.05), post hoc pairwise comparisons were conducted using Tukey’s Honestly Significant Difference (HSD) test.

For datasets violating normality assumptions, nonparametric alternatives were employed. The Kruskal-Wallis rank-sum test was first applied to assess global differences among groups, followed by Dunn’s test with Bonferroni adjustment for pairwise comparisons between specific treatments.

All statistical computations and graphical representations were generated using R statistical software (version 4.2.1, R Core Team 2023). The analysis workflow implemented specialized packages including ‘emmeans’ for estimated marginal means calculations and ‘agricolae’ for agricultural research-specific statistical procedures.

In the figures, groups not sharing the same letter designation (e.g., ‘a’ vs ‘b’) differ significantly (p < 0.05), while those with identical letters (e.g., ‘a’ and ‘a’) do not. Also, groups showing statistical difference are shown with asterisks, where one asterisk (*) indicates p < 0.05, two asterisks (**) represent p < 0.01, and three asterisks (***) signify p < 0.001. Comparisons marked with “ns” (not significant) indicate p ≥ 0.05.

Data for morphometric parameters are shown in tables 6, 7 Supplementary materials. The raw data used for gene expression analysis are not included in this article to maintain conciseness but are available from the corresponding author upon request.

## 3. Results

### 3.1. Morphometric parameters

Fresh weight measurements were not included in the statistical analysis due to their higher variability and non-normal distribution (Table 7, Supplementary material). Instead, we focused exclusively on root and shoot length as key morphometric parameters, as these traits exhibited a more stable, normally distributed pattern that better reflected the biological effects of the experimental treatments. This approach ensured greater reliability and interpretability of the statistical results.

The results of morphometric measurements are presented in Figures 1-6. Seven days after beginning of the experiment, almost no statistically significant difference was observed in the shoots and roots (Figure 1, 2). It is likely that no visible response was detectable at that time.

**Figure 1.**
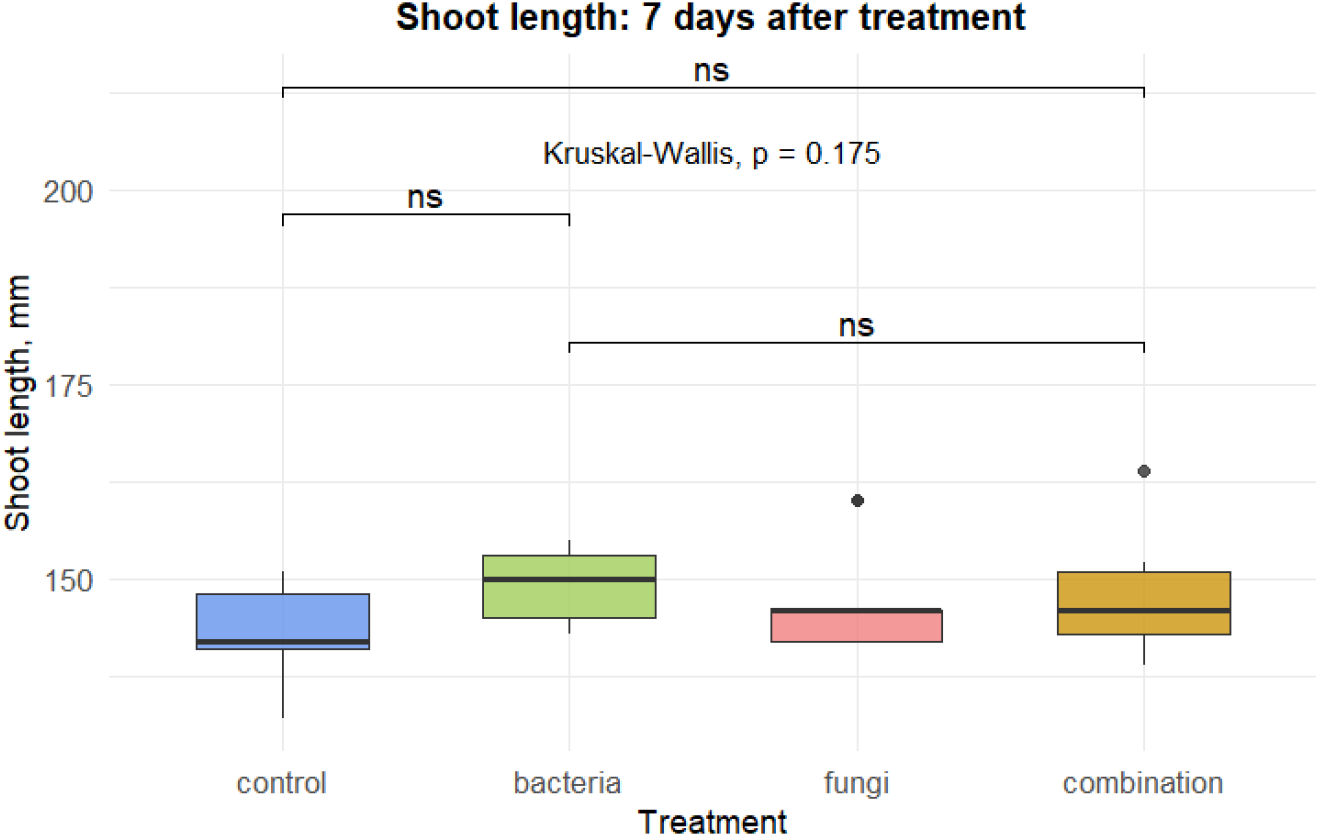
Length of the plant shoots 7 days after the beginning of the experiment

**Figure 2.**
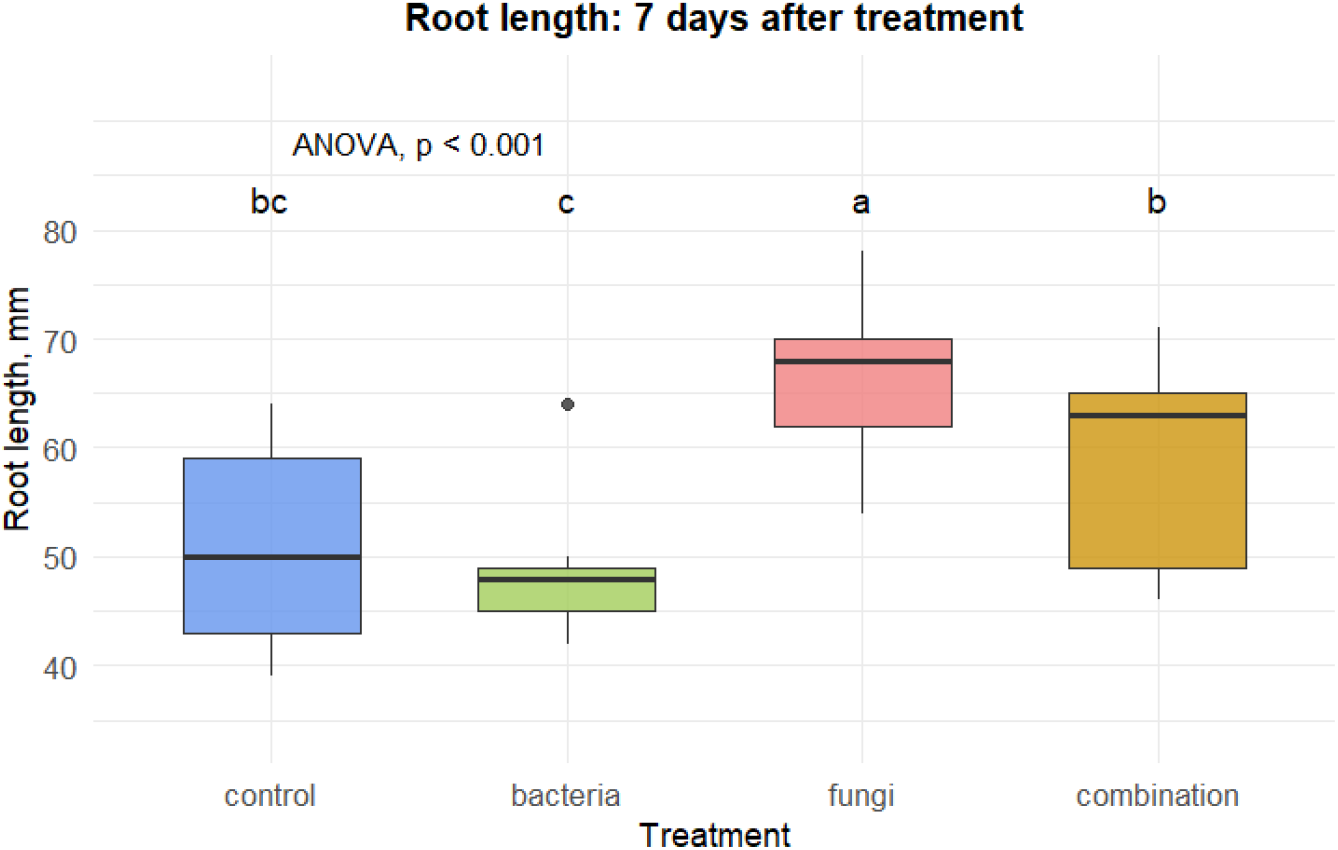
Length of the plant roots 7 days after the beginning of the experiment

After 14 days, statistically significant changes were observed between the control group and the groups treated with bacteria (Figure 3).

**Figure 3.**
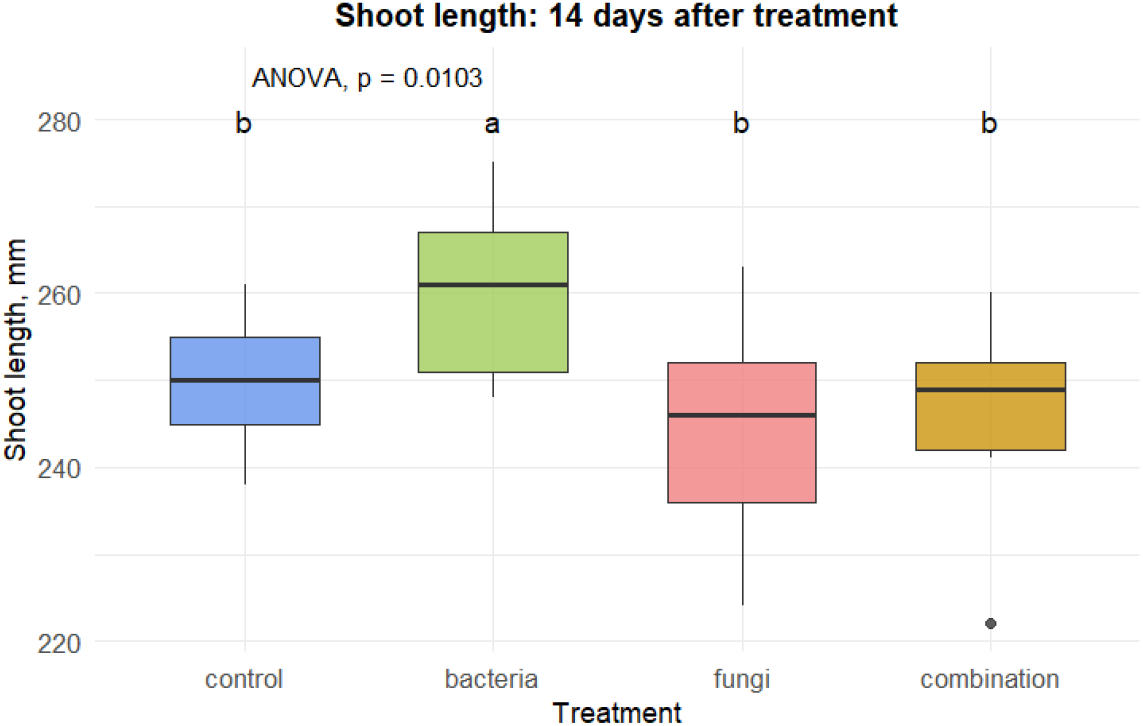
Length of the plant shoots 14 days after the beginning of the experiment

In the shoots, the bacteria-treated group showed 1,08 fold (8%) growth increase than the control one. In the roots, no significant difference was observed (Figure 4).

**Figure 4.**
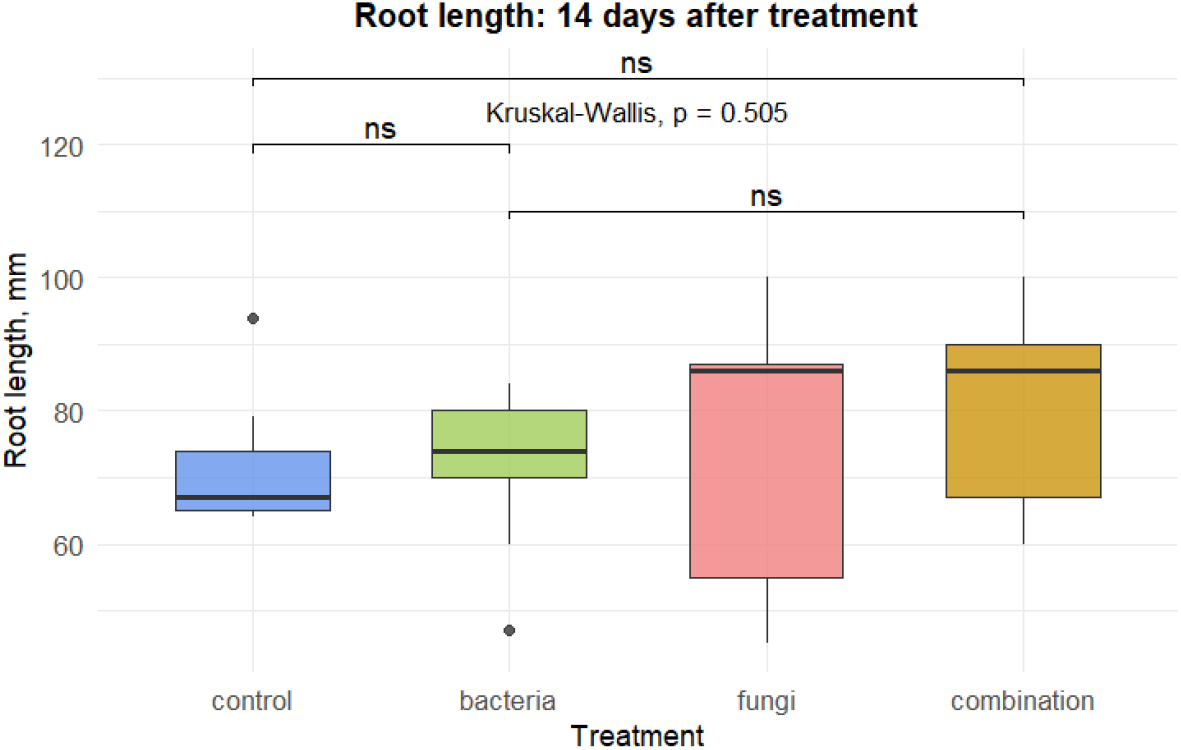
Length of the plant roots 14 days after the beginning of the experiment

After twenty-eight days after the start of the experiment, the shoot length of plants treated with bacteria showed statistically significant differences from the control group - thus, in the group treated with bacteria, seedling growth increased 1,08 fold (8%) compared to the control group (Figure 5).

**Figure 5.**
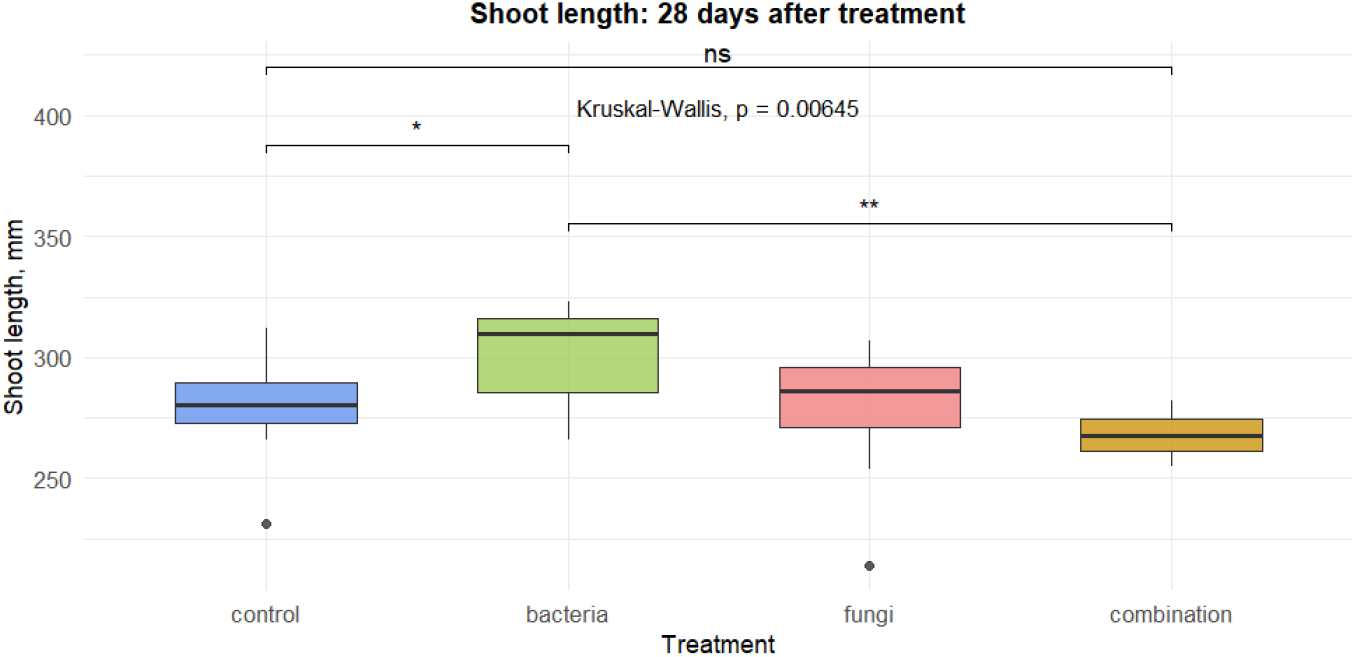
Length of the plant shoots 28 days after the beginning of the experiment

**Figure 6.**
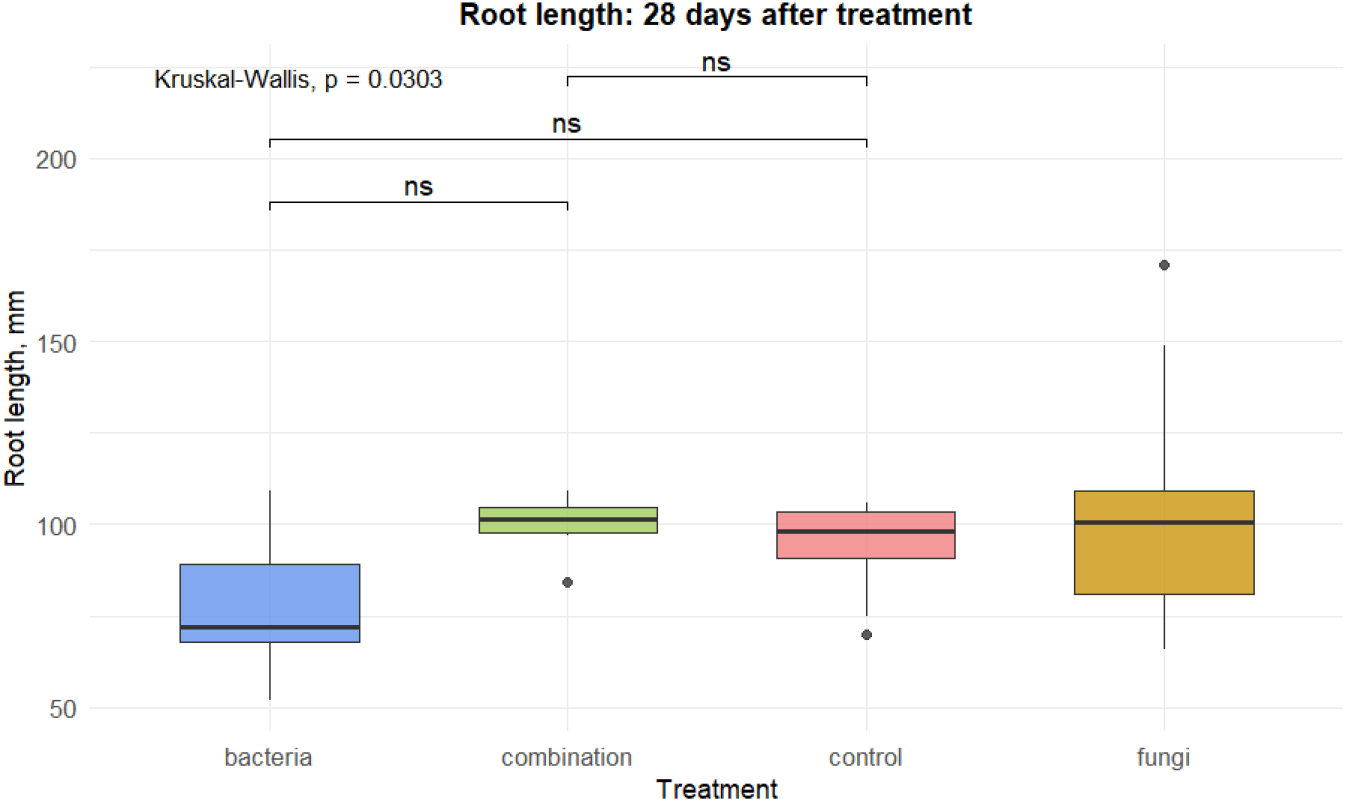
Length of the plant roots 28 days after the beginning of the experiment

Plants treated with bacteria were 1,08 (8%) times higher than plants treated with both bacteria and fungi. In the roots, bacteria-treated group showed 1,075 fold (7,5%) more intense growth than the control group (Figure 5).

No statistically significant difference in the root length in control and experimental groups was observed on this timepoint.

### 3.2. Expression levels of PR-genes in wheat

The results of differential expression measurements for *PR*-genes are presented in Figures 7– 12. Gene expression was measured in the lower part of shoots (approximately 2 cm from the root; referred to as ‘lower shoots’) and in the upper part, near the shoot apex (referred to as ‘upper/higher shoots’), and in roots.

**Figure 7.**
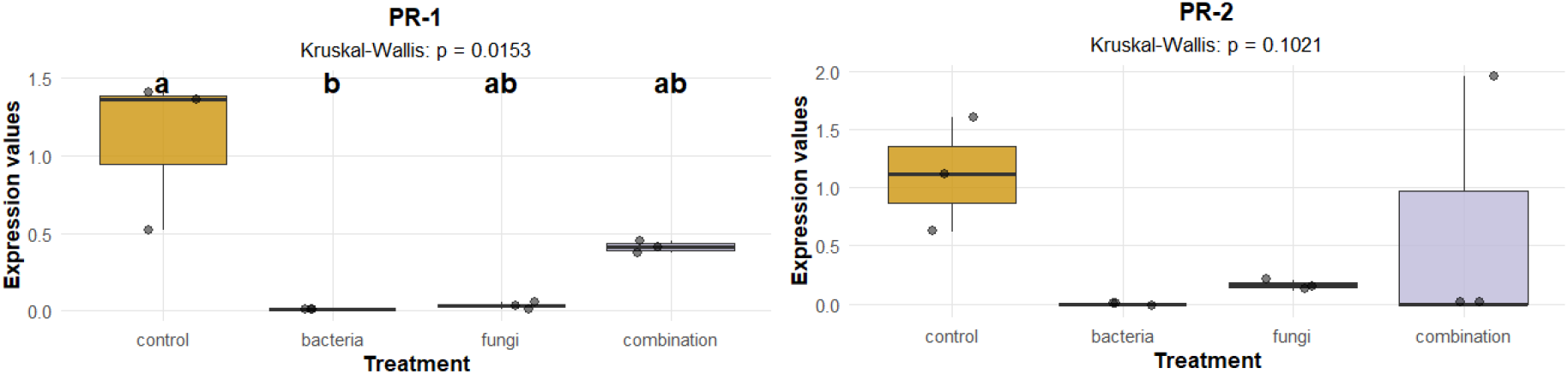
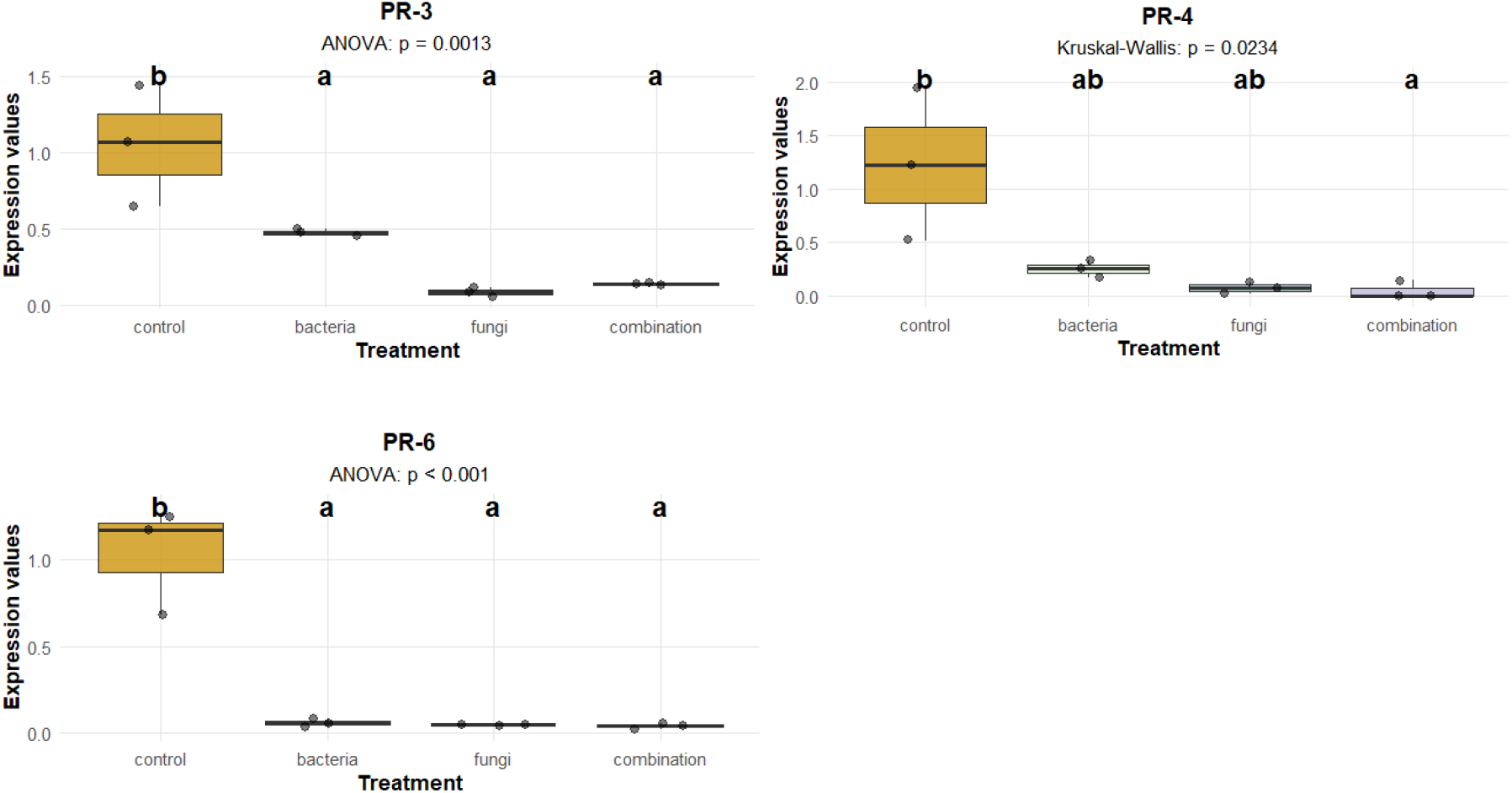
Expression of PR-genes in the upper shoots 7 days after the start of the experiment

Gene expression in the upper shoots of plants is presented in Figure 7. PR-3 was expressed in groups treated with fungi, bacteria, or fungi and bacteria simultaneously, with a statistically significant difference between the experimental and control groups.

*PR-4* followed a similar pattern: expression in the experimental groups was lower than in the control groups and close to zero. The mean value of *PR-2* expression in the group treated with fungi and bacteria simultaneously tends to be the highest among all genes in this case. The expression in control groups was approximately equal to one and had statistically significant differences with the experimental groups. *PR-6* expression in all experimental groups was near zero and differed statistically significantly from the control group. This effect may be due to delayed plant response to the exposure.

Seven days after the beginning of the experiment, decreased expression was observed in the lower shoots of plants in the experimental groups compared to control for all genes (Figure 8). *PR-1* expression in the groups treated with bacteria or with a combination of fungi and bacteria was lower than in control group. PR-2 was not expressed. *PR-3, PR-4* and *PR-6* expression levels for all experimental groups were lower than the control ones.

**Figure 8.**
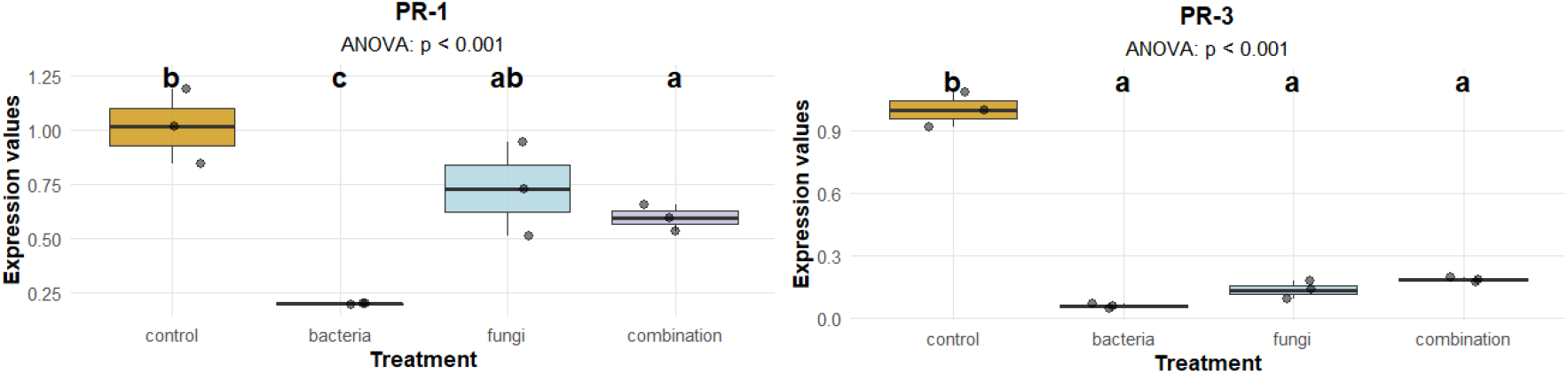
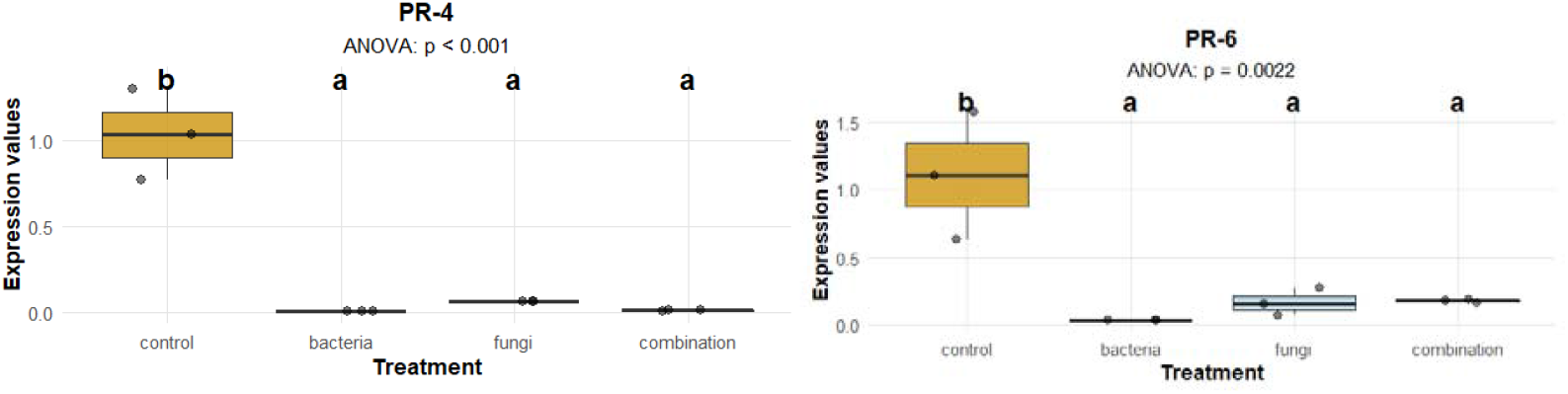
Expression of PR-genes in the lower shoots of plants 7 days after the start of the experiment

In contrast, in the roots after 7 days, the expression of *PR-1* and *PR-6* in experimental groups was mostly higher than in the control ones (Figure 9). *PR-1* expression in the bacteria-treated group was 16 times higher than in the control group.

**Figure 9.**
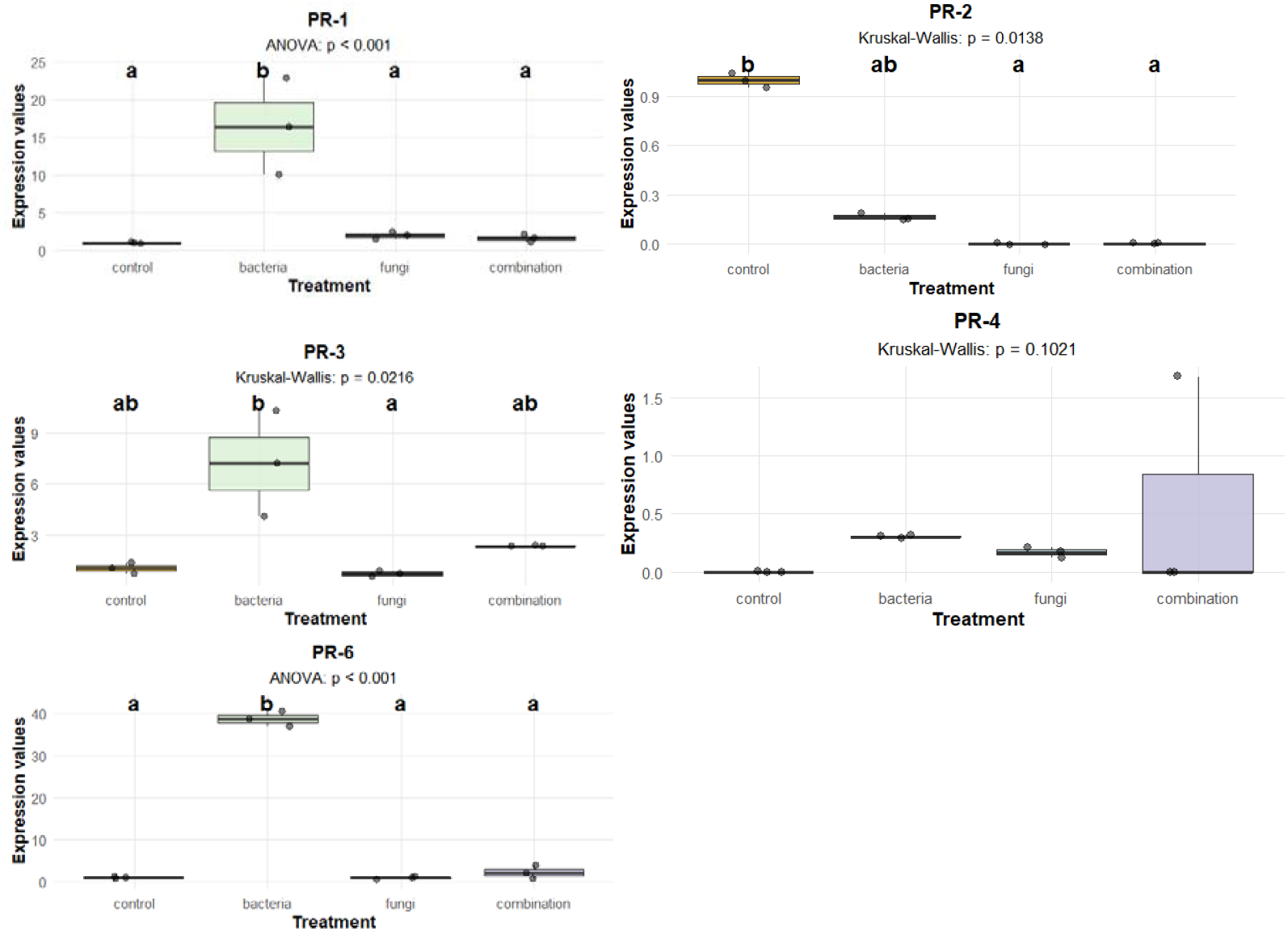
Expression of *PR*-genes in the root part of plants 7 days after the beginning of the experiment

The expression of *PR-2* in the control group higher than in groups treated with fungi and the combination of fungi and bacteria. Expression of *PR-3* in control group was 6 fold higher than this in fungi-treated group. *PR-6* expression in bacteria-treated was 38 times higher than in the control group.

Fourteen days after the start of the experiment, increased expression was observed in the upper shoots of the plants in the experimental groups compared to the control groups for *PR-2, PR-3, PR-4* and *PR-6* genes (Figure 10). *PR-2* expression in fungi-treated group was 4 times higher than in the control one. PR-3 expression in bacteria-treated group was 9 times higher than in the control group.

**Figure 10.**
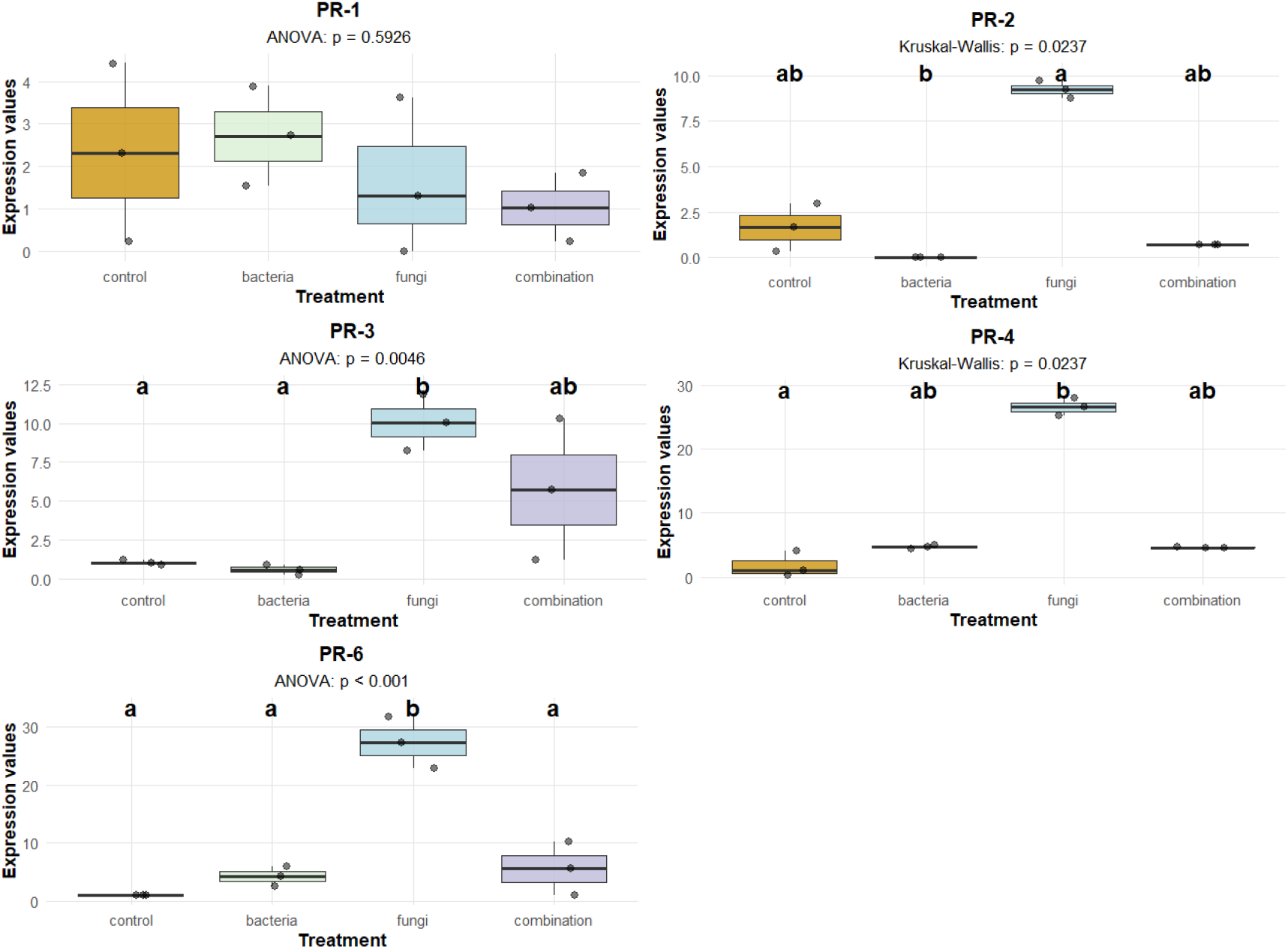
Expression of PR-genes in the upper shoots 14 days after the beginning of the experiment

*PR-4* expression in the group infected with fungi was 25-fold higher than in the control group. For *PR-6*, the group treated with fungi showed 25-fold higher expression.

The expression of the studied genes in the lower part of the plants after 14 days is shown in Figure 11. *PR-2* expression of when treated with bacteria was lower than in the control group. *PR-3* expression in fungi-treated group was 6 times higher than in the control group, and in fungi and bacteria treatment it was 2,5 times higher than in the control group.

**Figure 11.**
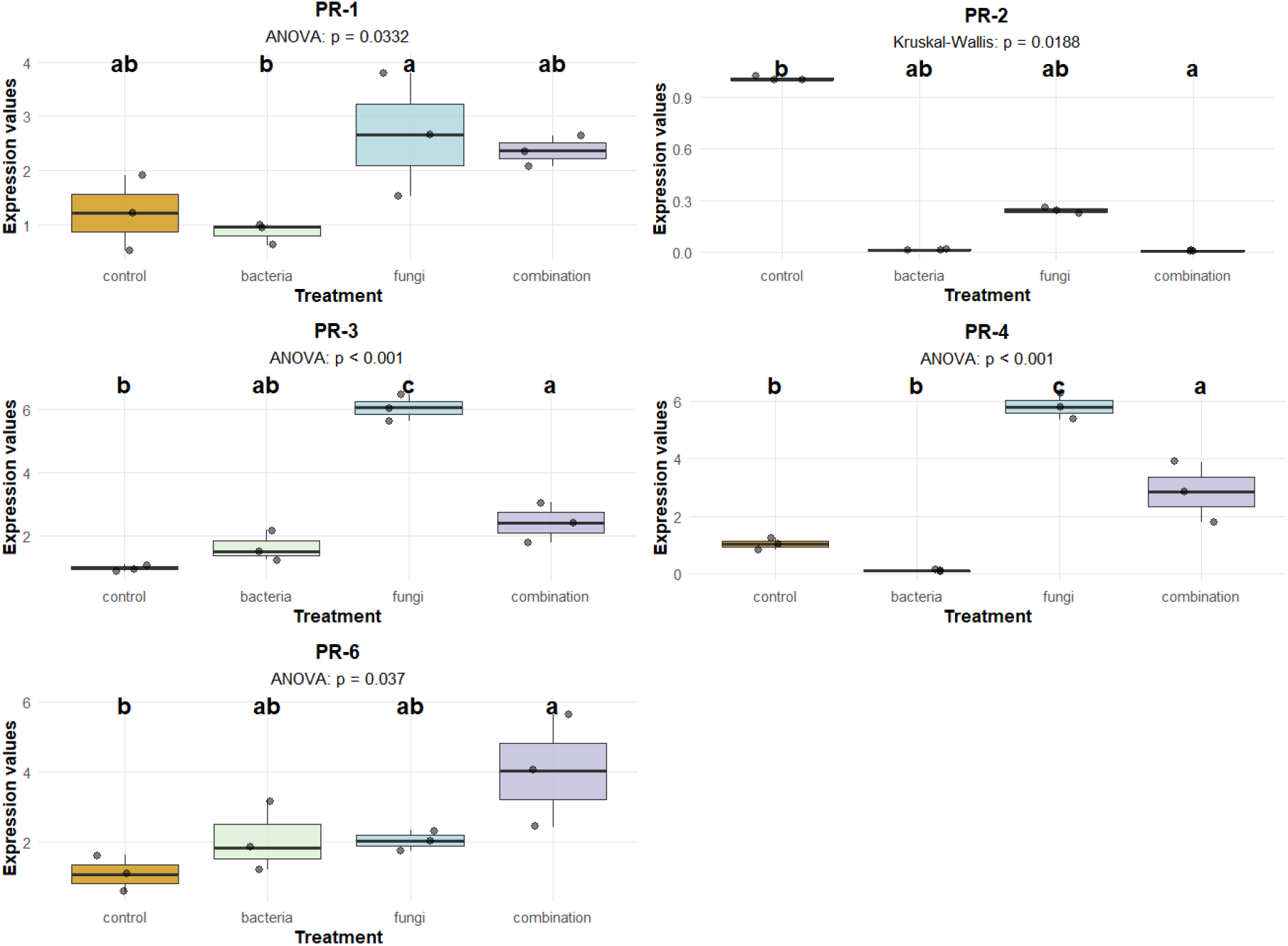
Expression of PR-genes in the lower shoots 14 days after the start of the experiment

*PR-4* expression in the fungi treatment was 5,5-fold higher than in the control group. The expression of PR-6 in the fungi-treated group was also 2-times higher, and in the bacteria and fungi treatments, it was 3,5 times higher than in control.

*PR*-gene expression in the roots was significantly different from that in the shoots (Figure 12). The expression of PR-1 when infected with bacteria was more than 400 times higher than in the control group. The expression of PR-2 and PR-4 was close to zero.

**Figure 12.**
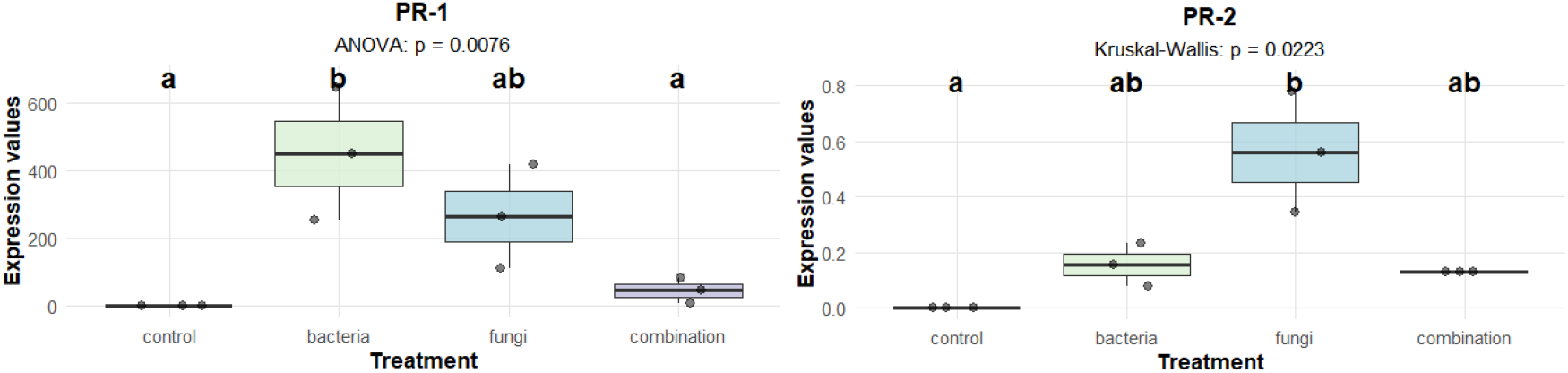
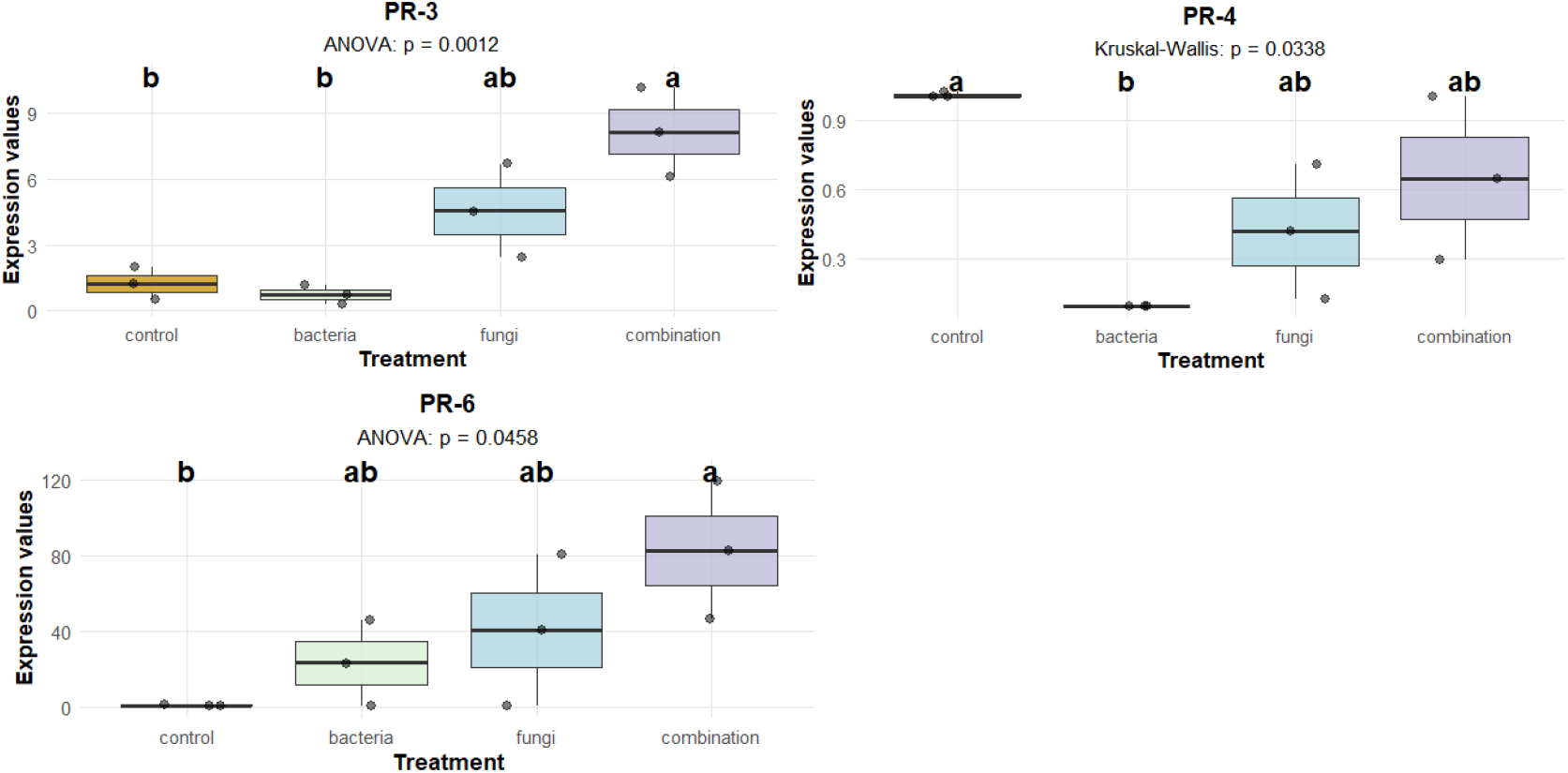
Expression of PR-genes in the root of plants 14 days after the start of the experiment

*PR-3* expression when treated with fungi and bacteria together was 6.5 times higher than in the control group. The expression of *PR-6* when treated with fungi and bacteria simultaneously was also 80 times higher than in the control group.

## 4. Discussion

The strains used in this study were described in detail in our previous works. Antagonistic interactions between soil bacteria of the genus *Bacillus* and target fungi were investigated, with the highest activity observed in bacteria isolated from soils where winter wheat was the preceding crop [13]. Genome analysis indicated that the antifungal effect of the strains is mainly due to non-ribosomal peptides, with enhanced expression of the corresponding genes during co-culture with fungi, which suggests a broad range of their biological activity [12].

### 4.1. Plant growth stimulation

It is known that *Bacillus* and *Paenibacillus* strains enhance wheat growth through several mechanisms. One of them is nitrogen fixation: certain strains, such as *Paenibacillus spp*., can fix atmospheric nitrogen, increasing soil nitrogen content, which is crucial for plant growth, and improving soil fertility [18]. It was also shown that *Bacillus spp*. enhance the uptake of essential macro-(nitrogen, phosphorus, potassium) and micronutrients (iron, zinc) by wheat. For instance, *Bacillus subtilis* increases mineral content in wheat grains and shoots, promoting overall plant development [19]. Furthermore, both *Bacillus* and *Paenibacillus* species produce phytohormones such as auxins (e.g., indole-3-acetic acid), cytokines, and gibberellins. These hormones regulate crucial processes in plants, such as cell proliferation, tissue growth and differentiation [17, 18]. Auxins are among the key regulators of plant growth, and auxin-dependent signaling processes influence lateral roots development and quantity [19, 20, 21]. Beyond metabolic regulation, *Bacillus* and *Paenibacillus* bacteria enhance plant resilience to abiotic stresses, such as salinity, drought, and heavy metals, by producing bioactive metabolites. For example, *Bacillus spp*. can alleviate the effects of salt stress by maintaining nutrient concentrations in stressed plants [25]. They also mitigate biotic stresses: *Bacillus* species can suppress soil-borne pathogens and improve the phytosanitary status of crops, protecting wheat from diseases and pests [26].

Gibberellic acid (GA3) accelerates photosynthesis, gas exchange, cell division and expansion, improves nutrient uptake (nitrogen, potassium, phosphorus) and promotes root, stem and leaf development. Gibberellins application increases fruit size and quality, delays senescence, reduces oxidative stress levels [27]. For example, orchid-associated *Fusarium proliferatum* synthesizes a significant amount of gibberellins [28]. Optimal cultivation conditions for *F. oxysporum* have been identified to enhance GA3 production during submerged fermentation process [29] and under solid state fermentation. GA3 extract improves growth and physiological parameters of tomatoes under salt stress, including plant height, chlorophyll, starch and proline content [30]. Finally, *Bacillus* strains can stimulate the rhizosphere microbial community, which in turn increases the availability of nutrients and enhances soil health, further benefiting wheat growth [31].

In the present study, differences between experimental and control groups in the length of the shoots were observed. In general, the growth dynamics can be described as follows: after 7 days, exposure to *Fusarium* stimulated root growth, which may be an adaptive mechanism to escape localized fungal pressure under stress or the action of gibberellin produced by the fungus. By day 14, growth stimulation of the aboveground part was observed with bacterial (1,08 fold) treatment, while the root systems leveled off in their development rates. This trend continued at day 28, with the bacterial consortium stimulating growth by 1,08 fold and the fungi-treated group also showing a 1,04 fold increase. Stimulation of shoot growth by bacteria indicates the ability of the selected strains to positively influence plant growth under normal conditions (in the absence of pathogen). No significant differences among groups in root growths on 28 day indicating that root growth effects are transient or compensated over time.

The findings are partially consistent with similar previous studies. For example, Saleemi et al. [32] demonstrated that applying bacteria isolated from wheat rhizosphere improved plant growth. Co-inoculation with four strains gave the best result (278 mm mm shoot length in the experimental group, 81 mm in the control group) after 8 weeks. In a study by Wang et al. [33] wheat exposed to *Fusarium graminearum* exhibited reduced shoot (7-9% less) and root (up to 13% less) lengths compared to the control group at 7, 14, and 21 days. Similar results [34] were reported for eight PGPR strains, including *Bacillus endophyticus* and two *Bacillus filamentosus strains*, and wheat “Saha 94”. After 90 days, the maximum shoot height of plants treated with *Micrococcus luteus* and *B. endophyticus* was 534 mm and 532 mm, respectively, compared to 376 mm in the control. *Bacillus pumilus* strain EU927414, in combination with another strain, increased shoot biomass by up to 28% under low-intensity salt stress [35]. In another study [7], it was found that *Bacillus megaterium, Arthrobacter chlorophenolicus*, and *Enterobacter sp*. increased plant height; after 60 days, control group growth was approximately 57 cm, while *B. megaterium* treatment resulted in approximately 64 cm growth, along with improved grain yield and micronutrient content.

The plant growth promotion by bacterial treatment in this research can likely be attributed to the previously described positive effects of PGPR on nutrient uptake and availability. The increased shoot growth in fungi-infected groups at days 14 and 28 could be due to signaling interactions between the fungi and the plant or other molecular mechanisms. The observed stimulation of root growth, though, is a novel finding. Typically, *Fusarium* exposure leads to neutral or negative effects on root growth. It was observed [36] that inoculation with *Fusarium culmorum* and Fusarium *pseudograminearum* not only reduced shoot size and biomass but also altered root system length, biomass and architecture.

### 4.2. Influence of on the PGPR expression of PR genes

PGPR can induce systemic acquired resistance (SAR) in plants, characterized by the upregulation of PR genes. These genes encode proteins involved in plant defense mechanisms against microbial challenges. Several studies have shown that specific PGPR strains can trigger the expression of *PR*-genes like *PR-1, PR-2*, and *PR-4*, associated with enhanced responses to Fusarium exposure in wheat. The activation of these genes is often linked to the production of signaling molecules such as salicylic acid, mediating plant defense responses [34, 35].

Besides direct induction, PGPR can also prime wheat plants for a more robust response to subsequent negative influence. This priming involves the enhancement of the plant’s innate immune response, making it more responsive to stress signals and pathogens. PGPR can modify the expression of genes related to stress responses and metabolism, leading to an increased readiness to activate PR-genes expression upon pathogen challenge [39].

In studies involving wheat varieties challenged with *Fusarium equiseti*, the expression of various *PR*-genes was significantly elevated in PGPR-treated plants compared to untreated controls, suggesting that PGPR enhance both growth and plant defense mechanisms against fungi [40]. In particular, PGPR application was shown to reduce the severity of *Fusarium* exposure in wheat. For example, *Bacillus amyloliquefaciens* exhibits antifungal properties and enhances the expression of *PR*-genes, leading to decreased disease incidence and increased crop yield [41].

### 4.3. Fusarium exposure affecting PR-genes

*Fusarium* treatment can significantly induce the expression of *PR*-genes in wheat, particularly in resistant genotypes. The magnitude of upregulation can range from several-fold to over a thousand-fold, depending on the specific *PR*-gene and the stage of the exposure. This induction is a crucial component of the plant’s defense response against *Fusarium* fungi.

*PR-1* genes are often used as markers for systemic acquired resistance in plants. *Fusarium* exposure can induce the expression of PR-1 genes in wheat, indicating defense responses activation [1]. *PR-2* genes encode β-1,3-glucanases, which hydrolyze fungal cell walls. Overexpression of β-1,3-glucanase genes in flax has been shown to enhance resistance against *Fusarium* species and participate in the regulation of interaction between infected and healthy cells [42]. *PR-3* genes encode chitinases, which can degrade fungal and bacterial cell walls. *PR-4* genes encode ribonucleases and thaumatin-like proteins with antifungal properties. *PR-6* genes encode proteinase inhibitors that can interfere with fungal enzymes [42].

The expression of these PR-genes is often upregulated in response to *Fusarium* treatment, contributing to plant defense [43]. The magnitude of expression level changes can vary depending on the wheat genotype (resistant or susceptible) and the stage of exposure. Resistant wheat genotypes tend to exhibit stronger *PR*-gene upregulation in response to *Fusarium* exposure compared to susceptible genotypes [43]. For example, genes that putatively encode PR-1, PR-2, PR-4 and PR-5 proteins have been identified in *Allium sativum cv. Ershuizao*, potentially involved in defense against Fusarium exposure. The expression of these genes, AsPR1c, d, g, k, AsPR2b, AsPR5a, c (in roots) and AsPR4a, b and AsPR2c (in stems and cloves) differed significantly between *Fusarium*-resistant and susceptible garlic lines, so it is suggested that the expression of these genes determines protection against *Fusarium* exposure [44].

*PR*-genes expression levels also tend to vary temporally and may increase several-fold during the early stages of exposure (12-48 hours post-infection) [45]. For instance, one study identified 360 upregulated and 219 downregulated *PR* genes common between 12 and 48 hours post-infection, with a more intense response at 48 hours [46].

In the current study, after 7 days in both upper and lower shoots, *PR*-gene expression (especially *PR-1, PR-3, PR-4, PR-6*) was generally downregulated in experimental groups compared to controls. This could reflect a delayed or suppressed local immune response, possibly due to fungal or beneficial microbe-mediated suppression of plant defenses to facilitate colonization. *Fusarium* suppresses plant immunity by secreting effector proteins like effectors FoSSP71 and FolSCP1, that reduce the expression of pathogenesis-related (PR) genes [44, 45].

In contrast, *PR-1* and *PR-6* were strongly upregulated in roots, particularly in the bacteria-treated group. This suggests that roots mount a defense response, possibly as a front-line barrier triggered by beneficial bacteria.

After 14 days gene expression in the shoots was generally upregulated in the bacteria-treated compared to the control. Expression in the roots varied, with *PR-3* and *PR-6* most intensely expressed in the fungi and bacteria-treated group. The strongly increased expression in this case is likely due to the effect of bacterial treatment, given that in the groups treated separately with bacteria or treated with fungi, the mean value is lower and closer to the control group. PGPR may have enhanced the immune response and increased expression of these two genes. A similar expression pattern was observed for PR-6 in the upper shoots after 14 days.

High upregulation of *PR-1* in Fusarium-treated and bacteria-treated roots, and strong induction of *PR-3* and *PR-6* when both microbes are present may indicate mutual suppression of *Fusarium* and antifungal bacteria.

This suggests that bacterial exposure increased *PR-6* expression, which is consistent with other studies reporting increased expression of immunity genes when exposed to the fungi and PGPR [49, 50]. However, in most experimental variants, the expression of *PR*-genes was higher either in the fungi treatment or in the bacteria treatment separately. Increased expression of immunity genes when infected with fungi separately is a normal plant response to the fungal exposure, and such a response in the experimental groups was detected after 14 days for *PR-2, PR-3, PR-4, PR-6* in the shoots and *PR-1* in the root part.

The strong upregulation of *PR*-genes in roots and later in shoots, particularly in bacteria-treated plants, is consistent with theory, where beneficial microbes prime the plant for enhanced defense upon subsequent fungal exposure.

It is worth noting that the greatest increase (hundreds-fold) was observed in the expression of the *PR*-1 gene in the roots. The 14-day time point possibly reflects the peak of the bacterial-induced response, when PR-1 expression reaches its maximum level. The biological significance of such high expression levels should be assessed in the context of the normal dynamic range and function of the gene. For defense-related genes such as *PR-1*, increases in expression levels of several hundredfold may reflect a normal biological response to microbial stimuli [51].

Studies have consistently shown that *PR-1* genes can exhibit tissue-specific expression patterns, with roots often showing different responses compared to shoots [52, 53]. This tissue specificity explains why dramatic increases in expression were observed in roots and not shoot tissues.

### 4.4. Effects of co-treatment of PGPB and Fusarium on PR-genes expression

Although simultaneous exposure to both PGPR and Fusarium fungi is generally expected to enhance the plant immune response, gene expression patterns vary considerably among plant species and across different studies. For example, in a study examining the combined effect of *Trichoderma atroviride* and *Bacillus subtilis* on the *Fusarium graminearum* spread in infected wheat, *PR-1* expression was hardly elevated compared to the control, and *PR-3* expression was only slightly elevated. *PR-4* expression increased more than 4-fold compared to the control when exposed to both cultures, and *PR-5* expression increased more than 2-fold [49].

Similar studies were conducted for other fungal pathogens: for example, the ability of *Bacillus subtilis Cohn* and *Bacillus thuringiensis Berliner* to induce systemic resistance of wheat plants to Septoria *nodorum Berk in vitro* was studied. Treatment with this strain was accompanied by expression of immunity genes *PR-1, PR-6* and *PR-9* [50].

However, not all immunity genes showed consistently increased expression under these conditions. *Trichoderma longibrachiatum* AD-1 cell-free culture filtrate effectively inhibited *Fusarium solani* growth and spore germination, reduced root rot in common bean, and enhanced plant survival. In this case, the filtrate upregulated defense-related genes (*PR1, PR2, PR3*, and *PR4*) in common bean, suggesting that this preparation might be a promising natural agent for controlling root rot diseases caused by *Fusarium spp*. [54]. This also indicates that not only the strain itself but also its cell-free metabolites might be effective against fusariosis.

All the abovementioned effects are summarized (Tables 3, 4, 5 in Supplementary materials) to enlighten the patterns in gene expression changes compared to different studies.

**Table 3.**
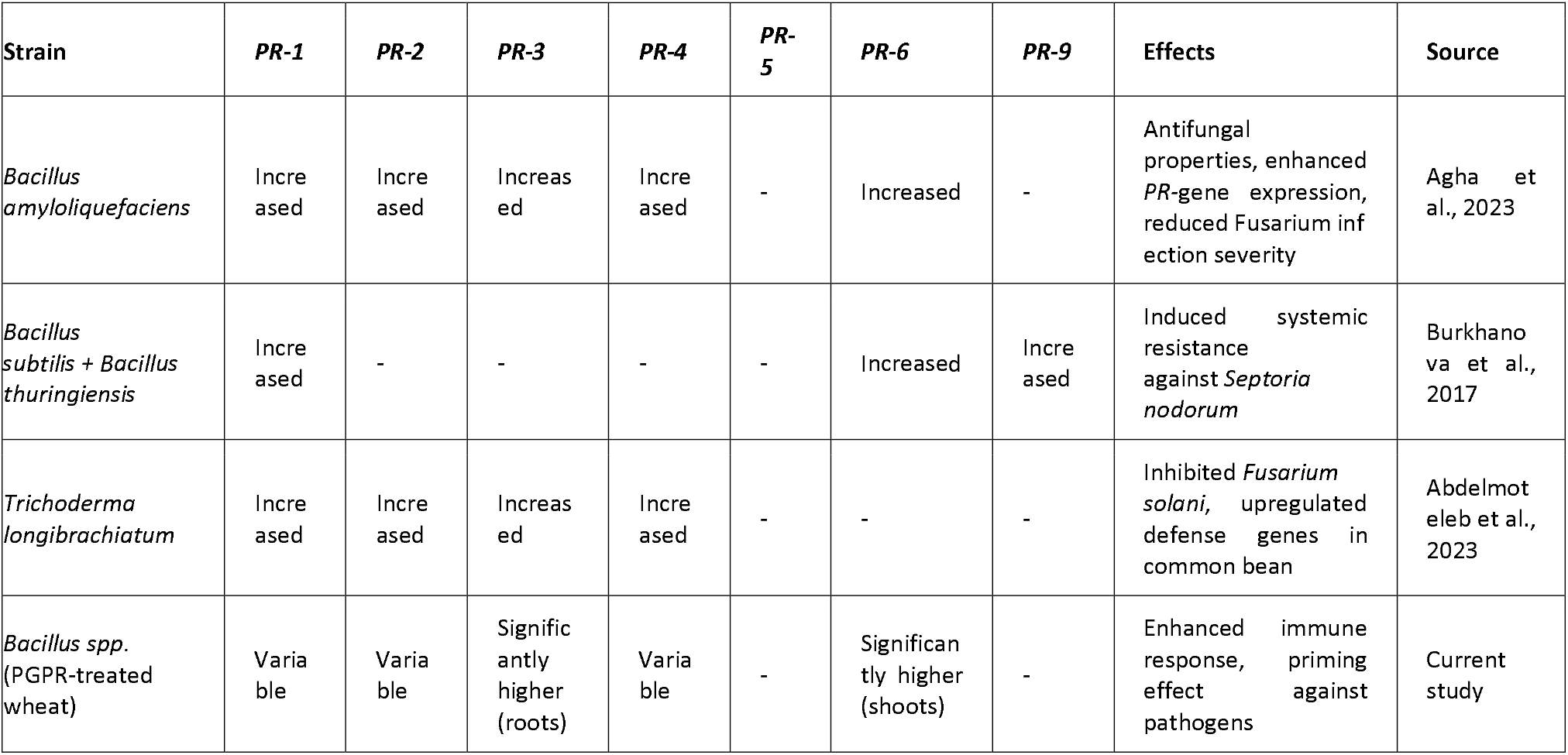
Summarized effects of bacterial treatment on *PR*-genes expression.

| Strain | *PR-1* | *PR-2* | *PR-3* | *PR-4* | *PR-5* | *PR-6* | *PR-9* | Effects | Source |
| --- | --- | --- | --- | --- | --- | --- | --- | --- | --- |
| *Bacillus amyloliquefaciens* | Increased | Increased | Increased | Increased | - | Increased | - | Antifungal properties, enhanced *PR-gene* expression, reduced *Fusarium* infection severity | Agha et al., 2023 |
| *Bacillus subtilis* + *Bacillus thuringiensis* | Increased | - | - | - | - | Increased | Increased | Induced systemic resistance against *Septoria nodorum* | Burkhanova et al., 2017 |
| *Trichoderma longibrachiatum* | Increased | Increased | Increased | Increased | - | - | - | Inhibited *Fusarium solani*, upregulated defense genes in common bean | Abdelmotaleb et al., 2023 |
| *Bacillus* spp. (PGPR-treated wheat) | Variable | Variable | Significantly higher (roots) | Variable | - | Significantly higher (shoots) | - | Enhanced immune response, priming effect against pathogens | Current study |

**Table 4.** Summarized effects of fungal treatment on *PR*-genes expression.

| Treatment | *PR-1* | *PR-2* | *PR-3* | *PR-4* | *PR-5* | *PR-6* | Effects | Source |
| --- | --- | --- | --- | --- | --- | --- | --- | --- |
| *Fusarium spp. exposure* | Significantly higher | Significantly higher | Significantly higher | Significantly higher | Significantly higher | Significantly higher | Strong induction of defense genes in resistant genotypes | Dmitriev et al., 2017 |
| *Fusarium graminearum* | Increased | Increased | Increased | Increased | Increased | Increased | Reduces shoot/root length but induces *PR*-genes | Wang et al., 2017 |
| *Fusarium culmorum* + *F. pseudograminearum* | - | - | Decreased root expression | - | - | Altered expression | Reduces root system length and biomass | Saad et al., 2023 |
| *Fusarium spp.* | Significantly higher | Significantly higher | Significantly higher | Significantly higher | Significantly higher | Significantly higher | Resistant genotypes show higher *PR*-gene upregulation than susceptible ones | Dmitriev et al., 2017; Anisimova et al., 2021 |
| *Fusarium spp.* | Generally lower | Generally lower | Generally lower | Generally lower | Generally lower | Generally lower | Infestation with phytopathogenic fungi leads to lower *PR*-genes expression | Current study |

**Table 5.** Summarized effects of fungal and bacterial treatment on *PR-genes* expression.

| Treatment | *PR-1* | *PR-2* | *PR-3* | *PR-4* | *PR-5* | *PR-6* | Effects | Source |
| --- | --- | --- | --- | --- | --- | --- | --- | --- |
| *Fusarium* + PGPR | Variable | Increased | Significantly higher | >4x increased | >2x increased | Increased | Combined effect stronger than single treatments | Hongyi et al., 2022 |
| *F. solani* + *T. longibrachiatum* filtrate | Increased | Increased | Increased | Increased | - | - | Filtrate inhibits fungal growth and upregulates defense genes | Abdelmotaleb et al., 2023 |
| *Bacillus subtilis* + *Fusarium graminearum* | - | - | Slight increase | >4x increased | >2x increased | - | Combined treatment increased PR-4 and PR-5 expression | Hongyi et al., 2022 |
| *Fusarium spp.* + *Bacillus* consortium | Increased | Variable | Significantly higher (roots) | Variable | - | Significantly higher (shoots) | Bacteria enhance plant response to fungal exposure | Current study |

**Table 6.** CV values for *cdc* and β -*tubulin* household genes’ Ct values in all the control groups.

| Samples | CDC | $\beta$ -tubulin |
| --- | --- | --- |
| upper shoots 7 days | 1,76 | 0,09 |
| lower shoots 7 days | 2,11 | 1,62 |
| roots 7 days | 1,78 | 7 |
| upper shoots 14 days | 2,65 | 2,17 |
| lower shoots 14 days | 0,9 | 3,16 |
| roots 14 days | 3,67 | 2,59 |

**Table 7.** Results of normality test for morphometric parameters.

| type | day | parameter | stat | p,value |
| --- | --- | --- | --- | --- |
| ctrl | 7 | height | 0,828170973 | 0,04602534 |
| bacteria | 7 | height | 0,911482137 | 0,326386382 |
| fungi | 7 | height | 0,971306149 | 0,905663517 |
| both | 7 | height | 0,849359082 | 0,073289364 |
| ctrl | 14 | height | 0,946718282 | 0,65410225 |
| bacteria | 14 | height | 0,911421481 | 0,325950739 |
| fungi | 14 | height | 0,958083374 | 0,778060342 |
| both | 14 | height | 0,904604022 | 0,279911847 |
| ctrl | 28 | height | 0,903036033 | 0,089901924 |
| bacteria | 28 | height | 0,957993759 | 0,746434042 |
| fungi | 28 | height | 0,948242284 | 0,291364626 |
| both | 28 | height | 0,855391436 | 0,173907028 |
| ctrl | 7 | roots_length | 0,9074536 | 0,298456872 |
| bacteria | 7 | roots_length | 0,787964181 | 0,014869464 |
| fungi | 7 | roots_length | 0,964929541 | 0,848230562 |
| both | 7 | roots_length | 0,877850108 | 0,149035227 |
| ctrl | 14 | roots_length | 0,756944956 | 0,006525403 |
| bacteria | 14 | roots_length | 0,877162049 | 0,146559599 |
| fungi | 14 | roots_length | 0,866629501 | 0,112816062 |
| both | 14 | roots_length | 0,916889371 | 0,36710904 |
| ctrl | 28 | roots_length | 0,861719483 | 0,020360608 |
| bacteria | 28 | roots_length | 0,946354543 | 0,597795777 |
| fungi | 28 | roots_length | 0,921018944 | 0,079805615 |
| both | 28 | roots_length | 0,908737274 | 0,428141328 |
| ctrl | 7 | roots_weight | 0,617277757 | 1,53E-04 |
| bacteria | 7 | roots_weight | 0,617277757 | 1,53E-04 |
| fungi | 7 | roots_weight | 0,891855945 | 0,20846745 |
| both | 7 | roots_weight | 0,898093212 | 0,241154194 |
| ctrl | 14 | roots_weight | 0,763013589 | 0,007670534 |
| bacteria | 14 | roots_weight | 0,805403233 | 0,023534082 |
| fungi | 14 | roots_weight | 0,805403233 | 0,023534082 |
| both | 14 | roots_weight | 0,53580985 | 1,69E-05 |
| ctrl | 28 | roots_weight | 0,816332277 | 0,004525345 |
| bacteria | 28 | roots_weight | 0,734022322 | 0,001287524 |
| fungi | 28 | roots_weight | 0,81530888 | 8,77E-04 |
| both | 28 | roots_weight | 0,496094403 | 2,07E-05 |
| ctrl | 7 | top_part | 0,955600813 | 0,751346751 |
| bacteria | 7 | top_part | 0,912856012 | 0,336378813 |
| fungi | 7 | top_part | 0,718381335 | 0,002325385 |
| both | 7 | top_part | 0,912102471 | 0,330868471 |
| ctrl | 14 | top_part | 0,958817137 | 0,785860303 |
| bacteria | 14 | top_part | 0,948965827 | 0,678709673 |
| fungi | 14 | top_part | 0,951702986 | 0,708764953 |
| both | 14 | top_part | 0,885060696 | 0,177381932 |
| ctrl | 28 | top_part | 0,90255693 | 0,088323846 |
| bacteria | 28 | top_part | 0,869966378 | 0,077444551 |
| fungi | 28 | top_part | 0,861776276 | 0,0055234 |
| both | 28 | top_part | 0,982159556 | 0,961775069 |
| ctrl | 7 | weight | 0,635903467 | 2,52E-04 |
| bacteria | 7 | weight | 0,830227736 | 0,044918187 |
| fungi | 7 | weight | 0,854529988 | 0,083566693 |
| both | 7 | weight | 0,953722852 | 0,730895584 |
| ctrl | 14 | weight | 0,806852829 | 0,024445822 |
| bacteria | 14 | weight | 0,941222893 | 0,594832282 |
| fungi | 14 | weight | 0,943611849 | 0,620391049 |
| both | 14 | weight | 0,71275035 | 0,00199917 |
| ctrl | 28 | weight | 0,923072351 | 0,18899628 |
| bacteria | 28 | weight | 0,979348182 | 0,962429399 |
| fungi | 28 | weight | 0,956376845 | 0,419686208 |
| both | 28 | weight | 0,894333514 | 0,341519158 |

### 4.5. Possible mechanisms of PGPB action

The interaction between PGPR and wheat involves complex signaling pathways. The upregulation of *PR*-genes in response to PGPR treatment is often associated with the activation of NPR1 (Non-expressor of *PR*-genes 1), a key regulator in the SAR pathway. This indicates that PGPR can modulate the expression of *PR*-genes through both direct and indirect pathways, enhancing the plant’s ability to resist *Fusarium infections* [55].

The metabolites responsible for these effects have not been identified in every such study. There are suggestions that non-ribosomal peptides (NRPs) may be such metabolites, acting at sub-inhibitory concentrations. At high concentrations, they exhibit antifungal properties, and retard the spread of *Fusarium*. Examples include surfactin, iturin, and fengycin [56], fusaricidin and polymyxin [51]. However, these compounds are also likely capable of promoting systemic resistance. Several bacterial lipopeptides, particularly those derived from the *Bacillus* bacteria, have been identified as effective inducers of systemic resistance in plants. Notably, surfactin and fengycin, both produced by *Bacillus subtilis*, elicit induced systemic resistance (ISR) responses, enhancing plant defenses against a variety of pathogens, including fungi and bacteria [52, 53].

Surfactin has been shown to trigger induced systemic resistance in various plant species such as beans and tomatoes, thereby strengthening their defense mechanisms against pathogens without causing significant phytotoxic effects [52, 54, 55]. Fengycin also effectively induces ISR, stimulating plant defense pathways and contributing to resistance against biotic stress [58].

The induction of ISR by these lipopeptides presumably involves “priming effect”, where lipopeptides prepare the plant to respond more rapidly and effectively to subsequent pathogen attacks without requiring extensive transcriptional changes beforehand. The ISR response is often mediated through various signaling pathways, including those involving salicylic acid, jasmonic acid, and ethylene, which are crucial for activating defense genes in plants [35, 54].

According to genomic data, these strains are able to produce various NRPS, including surfactin and fengycin. The study identified 10 Bacillus and *Paenibacillus* strains with antifungal activity linked to NRPS genes. Highly antagonistic strains exhibited strong NRPS expression, while weaker strains showed minimal activity. Notably, *Bacillus velezensis* strains carried surfactin and fengycin synthetase genes, while *Paenibacillus polymyxa* strains possessed fusaricidin and polymyxin synthetases. NRPS expression varied depending on the *Fusarium* species present, suggesting strain-specific antifungal responses. For example, fusaricidin production increased against *Fusarium oxysporum*, while fengycin was more active against *Fusarium graminearum*. These findings highlight NRPS-derived peptides as key antifungal agents, with potential for targeted biocontrol strategies [12].

To sum up, in current study an enhanced plant immune response was observed in the shoots of wheat plants when infected with fungi after 14 days, and this immune response was less pronounced when plants were exposed additionally to PGPR potential bacteria.

## 5. Conclusion

The study demonstrated that treatment of winter wheat with *Fusarium* and PGPR of the genera *Bacillus and Paenibacillus* affected the expression of plant immunity genes and morphometric parameters of plants. Plants treated with bacteria showed no initial difference in aboveground growth, but (fungus-infected) had longer roots compared to control groups. After 14 and 28 days, treated plants grew taller and developed stronger immune responses. An enhanced plant immune response was observed in the shoots when exposed to fungi, while the immune response became less pronounced when exposed additionally to PGPR potential bacteria. Therefore, these bacterial strains stimulate plant immunity and may help wheat resist fungal exposure, suggesting their potential use, in various combinations, as a biopreparation to improve winter wheat growth.

## 6. Author Contributions Statement

Conceptualization: V. A. Chistyakov, N.G. Vasilchenko; methodology: N.G. Vasilchenko, K. Mekhantseva, A.V. Usatov; formal analysis: K. Mekhantseva, N.G. Vasilchenko; investigation: K. Mekhantseva, N.G. Vasilchenko; resources: V.A. Chistyakov, E.V. Prazdnova, A.V. Usatov; data curation: K. Mekhantseva, E.V. Prazdnova; writing—original draft preparation: K. Mekhantseva; writing—review and editing: all authors; visualization: K. Mekhantseva; supervision: V.A. Chistyakov, E.V. Prazdnova, A.V. Usatov; project administration: V.A. Chistyakov, E.V. Prazdnova, A.V. Usatov; funding acquisition: V.A. Chistyakov. All authors have read and agreed to the published version of the manuscript.

## 7. Declaration of interest statement

The authors report there are no competing interests to declare.

During the preparation of this work the authors used “Gemini” in order to check correct translation of specific terms in context. After using this tool, the authors reviewed and edited the content as needed and take full responsibility for the content of the published article.

## 8. Data availability statement

The raw data supporting the findings of this study (gene expression Ct values) are available from the corresponding author, [Kamilla Mekhantseva], upon reasonable request. All other data generated or analysed during this study are included in this published article and its supplementary information files.

## 11. Appendices

